# Reward-Bases: Dopaminergic Mechanisms for Adaptive Acquisition of Multiple Reward Types

**DOI:** 10.1101/2023.05.09.540067

**Authors:** Beren Millidge, Yuhang Song, Armin Lak, Mark E. Walton, Rafal Bogacz

## Abstract

Animals can adapt their preferences for different types for reward according to physiological state, such as hunger or thirst. To describe this ability, we propose a simple extension of temporal difference model that learns multiple values of each state according to different reward dimensions such as food or water. By weighting these learned values according to the current needs, behaviour may be flexibly adapted to present demands. Our model predicts that different dopamine neurons should be selective for different reward dimensions. We reanalysed data from primate dopamine neurons and observed that in addition to subjective value, dopamine neurons encode a gradient of reward dimensions; some neurons respond most to food rewards while the others respond more to fluids. Moreover, our model reproduces instant generalization to new physiological state seen in dopamine responses and in behaviour. Our results demonstrate how simple neural circuit can flexibly optimize behaviour according to animals’ needs.

## Introduction

The dominant theory of dopamine function in the basal ganglia system is model-free reinforcement learning (RL) where the dopamine neurons in the Ventral Tegmental Area (VTA) encode reward prediction errors which are used to modulate plasticity at cortico-striatal synapses so as to learn a value function of cortical states (Houk & Adams, 1995; Montague, Dayan, & Sejnowski, 1996; Schultz, Dayan, & Montague, 1997). This theory is supported by large amount of experimental data showing that dopamine neurons produce patterns of activity consistent with reward prediction error (Cohen, Haesler, Vong, Lowell, & Uchida, 2012; D’Ardenne, McClure, Nystrom, & Cohen, 2008; Romo & Schultz, 1990; Tobler, Fiorillo, & Schultz, 2005) and modulate cortico-striatal plasticity in the directions predicted by the theory (Reynolds, Hyland, & Wickens, 2001; Shen, Flajolet, Greengard, & Surmeier, 2008). However, a key assumption of this model is that the amount of reward from different outcomes is fixed, while for biological organisms the ‘reward function’ can fluctuate over time depending on physiological state – e.g. food is rewarding when hungry but not when satiated.

There is substantial evidence that animals can flexibly adapt their behaviour in the face of changes to physiological needs, and they can instantly adapt their preferences without having to experience the rewards in the new physiological state. A compelling demonstrations of this capability come from a series of experiments (Robinson & Berridge, 2013) involving rats which learn to associate two levers with receiving either pleasant sugar juice or extremely unpleasant salt water. When they were placed in a physiological state which mimics the brain chemistry of salt deprivation, which they have never experienced before, they immediately approach the lever which is associated with the salt water. This cannot be explained through standard RL algorithms such as TD learning which require an experience of reward to update the value function estimate. TD learning would predict that the salt-deprived animals would still avoid the salty lever until the salt solution is delivered which would give them an unexpectedly positive reward, and would slowly induce them to approach that lever more and more often until it becomes the primary lever to be approached. Beyond this classic demonstration, there is a wide range of literature arguing that Pavlovian learned associations appear to be dynamically responsive not just to experienced rewards but to internal physiological states (DiFeliceantonio, Mabrouk, Kennedy, & Berridge, 2012; Krashes et al., 2009; Peciña & Berridge, 2013; Smith, Berridge, & Aldridge, 2011).

There have been several explanations proposed for this instant reward revaluation capability in the literature. It has been proposed that the ability to achieve instant reward revaluation is due to model-based planning (Daw, Niv, & Dayan, 2005; Dayan & Berridge, 2014). Another approach is to use successor representations (Dayan, 1993) which represent a ‘successor matrix’ of discounted state occupancies allowing value functions to be estimated for any given reward function. However, these approaches are computationally complex and thought to be implemented in the cortex (Gläscher, Daw, Dayan, & O’Doherty, 2010; Russek, Momennejad, Botvinick, Gershman, & Daw, 2021), while controlling behaviour based on physiological state is so fundamental for survival that is present in animals without cortex, such as drosophila where it is mediated by dopamine neurons (Senapati et al., 2019). In vertebrates dopamine neurons innervate the basal ganglia system, which is a core component in reward-based action selection (Daw & Dayan, 2014; Dayan & Balleine, 2002; Schultz, 2016; Schultz et al., 1997; Sesack & Grace, 2010; Tanaka et al., 2016), and is linked to brain systems controlling physiological state such as the hypothalamus (Godfrey & Borgland, 2019; Kelley, Baldo, Pratt, & Will, 2005; Morales & Berridge, 2020; O’Connor et al., 2015). The ability of basal ganglia for controlling behavior based on physiological state is consistent with observations that the dopamine release depends on physiological state (Cone et al., 2016; Papageorgiou, Baudonnat, Cucca, & Walton, 2016), as well as that dorsomedial striatal lesions abolish flexible reward devaluation behaviour (Yin, Ostlund, Knowlton, & Balleine, 2005). Moreover, there is evidence that the basal ganglia model-free system may be relied on *more* than the model-based system when physiological needs are pressing (van Swieten, Bogacz, & Manohar, 2021). There is thus a fundamental difficulty with the standard theory: the basal ganglia needs to flexibly adapt to changing reward functions to effectively control behaviour, yet TD learning, the algorithm it supposedly implements, simply cannot do this. In this paper, we propose a simple extension to TD learning that allows for instant generalization to changing reward functions. Specifically, we demonstrate that if we interpret the reward function as a linear combination of *reward basis vectors* and then learn a separate value function for each reward basis using standard TD learning, then we can instantly compute the value function of any reward function in the span of the reward basis vectors. Moreover, this algorithm can be straightforwardly implemented in neural circuitry by simply parallelizing the circuits proposed by Schultz (1998). Our model predicts that different dopamine neurons should encode prediction errors connected with different reward dimensions, and we verify this prediction by reanalysing existing data from primate dopamine neurons (Lak, Stauffer, & Schultz, 2014). Additionally, we demonstrate that our model can reproduce instant generalization to new physiological state seen in dopamine responses (Papageorgiou et al., 2016) and behaviour (Robinson & Berridge, 2013), and achieves similar performance as successor representations (Dayan, 1993) at RL tasks while using much less memory.

## Results

### Reward Bases model

Following standard RL, we denote the current state of an agent by *x*. This state may correspond to animal’s location in space or a stimulus presented during an experiment. We also denote the reward received in the current state by *r*(*x*). As in standard TD model, we assume that the goal of the learning process it to estimate the value function of states defined as:

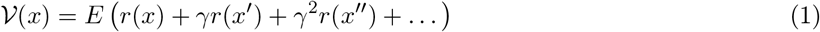

In the above equation, *E* denotes an expectation, *x′* denotes a state in which the agent finds itself in the next time step, and *γ* denotes discount rate expressing how much a reward in the next time step is worth to the agent relative to the same reward in the current time step. Thus the value function 𝒱 (*x*) expresses how much reward is expected in state *x* immediately and into the future, while considering that reward in the future is worth less than the reward now.

The key novel assumption of our model is that the reward can be decomposed into a linear combination of component rewards, or *reward bases*:

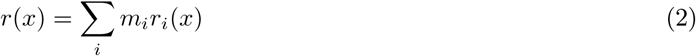

where *r*_*i*_(*x*) denotes an individual reward basis, and *m*_*i*_ is a motivational drive parameter for each reward basis. For instance, if *r*_*i*_(*x*) is a reward basis which represents food, then *m*_*i*_ would reflect the degree of hunger – i.e. how much food is valued by the agent. We assume that the basal ganglia system represents those dimension of reward that are most important for survival (the evidence shown later suggests they include food and water, but the exact range of dimensions will need to be identified by future studies). By substituting the definition of reward bases (Equation 2) into that of the value function (Equation 1), we can see that the value function decomposes into a weighted sum of component value functions,

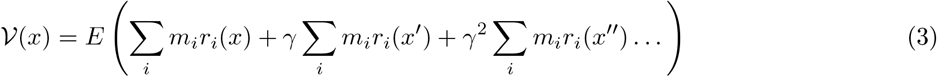

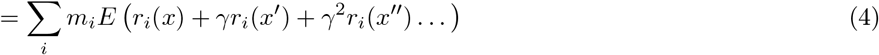

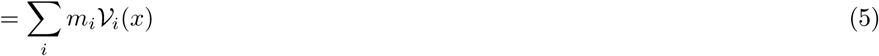

where we define 𝒱 _*i*_(*x*) = *E(r*_*i*_(*x*) + *γr*_*i*_(*x*^*′*^) + *γ*^2^*r*_*i*_(*x*^*′′*^) …) as a *value basis* since it is the value function of one specific reward basis. All of these value bases can be learnt using an analogous rule as in the TD learning, but using the reward basis as the reward instead of the full reward,

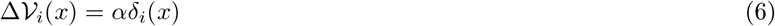

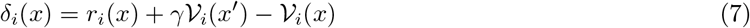

where *α* denotes the learning rate, and *δ*_*i*_(*x*) is a prediction error associated with reward basis *i*, defined analogously as in standard TD learning (cf. Equations 8 and 9 in Methods) but for just a single reward dimension. Given a set of value bases, and a set of coefficients {*m*_*i*_}, we can instantly compute any value function spanned by the reward bases. This enables instant generalization of the optimal value function for *any* reward function expressable as a linear combination of the reward bases. Equations 6 and 7 describe learning in the fundamental version of the Reward Bases model, and later in the paper it will be shown how these rules can be further refined to capture specific experimental data, but for simplicity we start with analysing this version.

To build intuition for how the Reward Bases model functions, in Figure 1 we present a visualization of the value functions learned by a TD and Reward Bases agents in a simple spacial navigation task. The agents must navigate around a 6 *×* 6 grid where they can obtain rewards in three locations (indicated in green or yellow in the top row). We assume that these three rewards are of different types, and activate separate reward bases (Figure 1B top). The TD agent received a total reward defined as 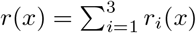 (Figure 1A top).The value function learnt by the TD and Reward Bases agents are shown in the middle row of Figure 1. Using the set of value bases in Figure 1B and equal weighting coefficients of 1, the Reward Bases agent can exactly reconstruct the total value function of the TD agent (cf. Figure 1A middle and Figure 1B middle right). Importantly, the Reward Bases agent can also instantly generalize to other reward preferences. For instance, when only the first reward becomes full valuable, the second only keeps half of its value, and the third is not valuable, the Reward Bases agent can choose its actions according to the appropriate value function shown in Figure 1B bottom.

**Figure 1:**
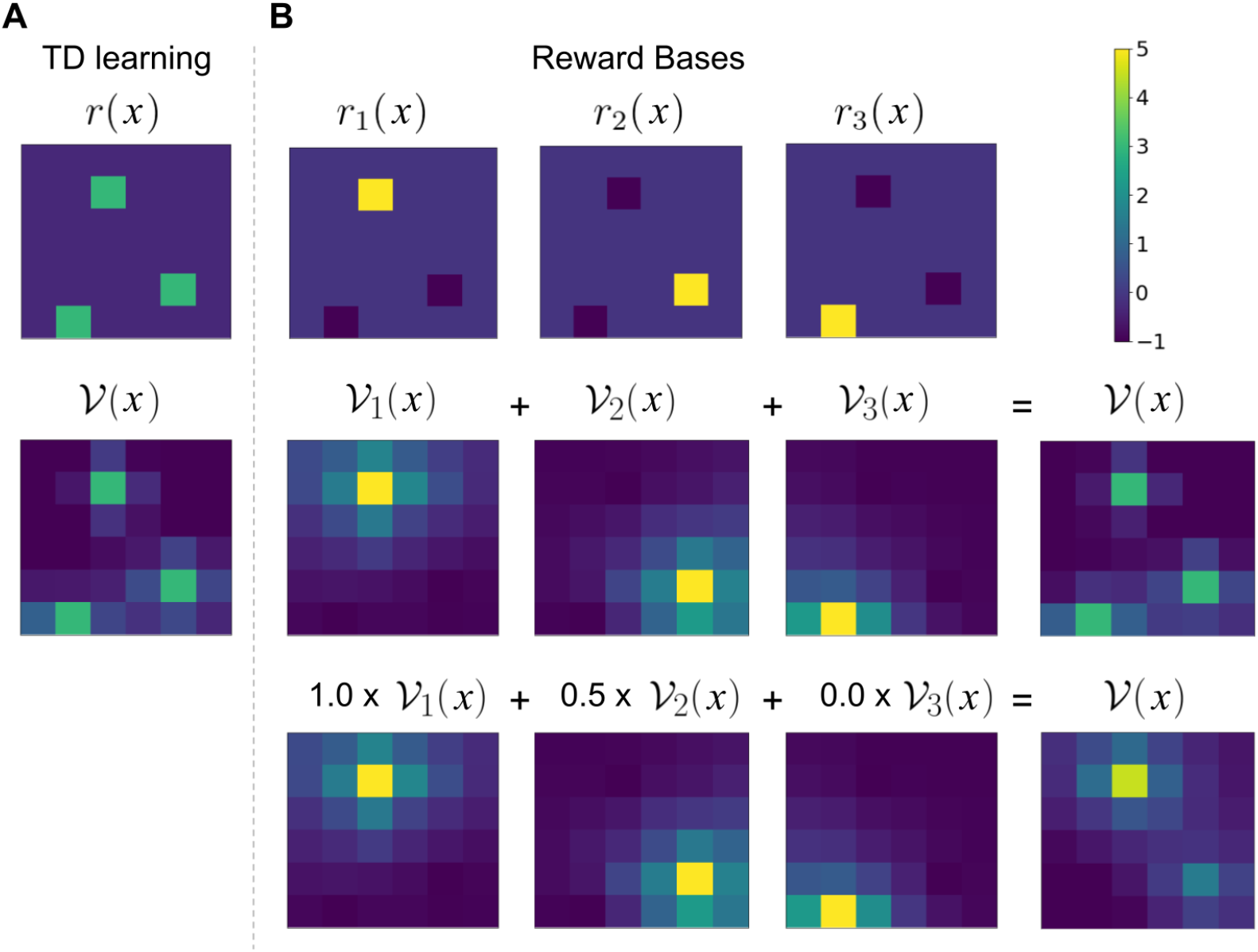
Visualization of the value functions learned by the models. A) TD model. B) Reward Bases model. The task consisted of a 6 × 6 grid with three items. The Reward Bases agent used a separate reward basis for each object, giving it +5 for that object, −1 for the other objects and −0.1 for all other squares. The value functions displayed are obtained by the agents exploring the environment for 1000 steps with a random action policy, learning rate *α* = 0.1 and discount factor *γ* = 0.99.

The Reward Bases model also possesses a potentially straightforward implementation in the circuitry of the basal ganglia. This is achieved as a simple modification of the original circuit model of Schultz (1998) illustrated in Figure 2A. In this model, dopamine neurons in the VTA receive two inputs – the incoming reward and the temporal derivative of the value function. To compute the reward prediction error, the dopamine neurons simply need to add these two inputs. These reward prediction errors can then be sent to the ventral striatum where they are used to update synaptic weights encoding the value function. The mapping of the Reward Bases algorithm on a neural circuit is shown in Figure 2B, and the only change is that instead of assuming a single homogenous population of dopamine neurons responding to a global reward signal, we instead assume that neurons encoding values and prediction errors are parcellated into groups which each responding to a single reward basis. This reward basis parcel is simply the same circuit as original model in miniature in that dopamine neurons simply receive both the reward for that basis, and the temporal derivative of the value function for that basis and compute the reward prediction error for that basis, which is then sent specifically to the region which requires it. A set of parallel units represents the value bases, from which the ultimate value function can be computed through a simple linear combination with the weighting coefficients (cf. Khamassi, Lachèze, Girard, Berthoz, and Guillot (2005)). Hence there need to also exist inputs to each reward basis parcel from neurons encoding corresponding physiological need, but they are not included in the diagram for simplicity. The fundamental prediction of the Reward Bases model is that individual dopamine neurons should signal reward prediction errors for a specific reward basis, and below we compare this prediction with experimental data.

**Figure 2:**
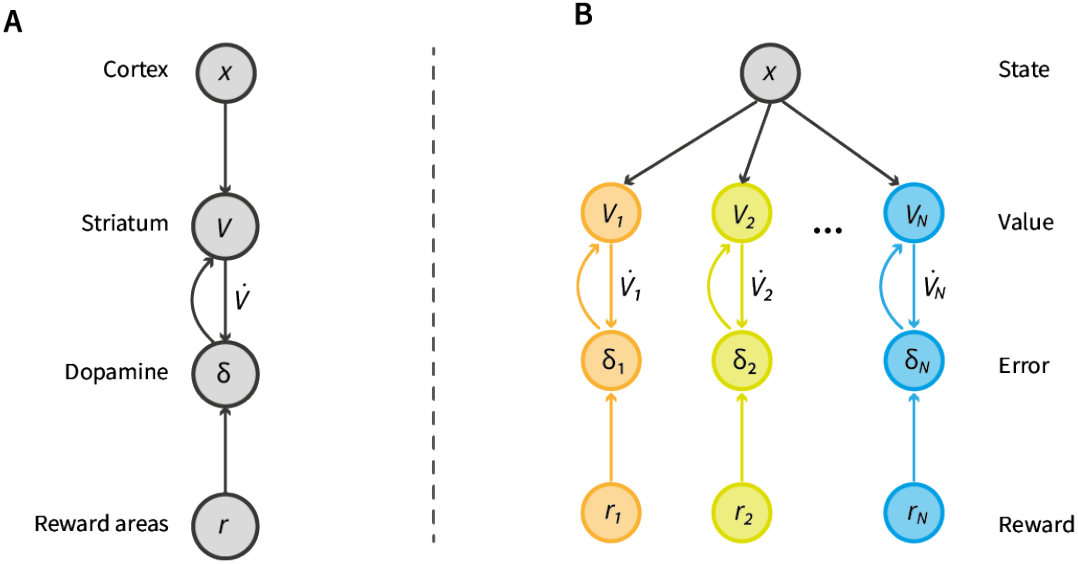
Possible neural implementations of TD learning and the Reward Bases model. **A**: Standard TD learning model. Dopamine neurons receive reward input and compute a reward prediction error. This reward prediction error is projected to the striatum where it modulates plasticity of the corticostriatal synapses that learn a value function. **B**: Reward Bases model. This requires parallel neural populations computing prediction errors and ‘value function’ neurons with the same connectivity patterns as the standard model.

### Different dopamine neurons are selective for different reward dimensions

Given that a core prediction of the Reward Bases model is that dopamine neurons in the VTA should be differentially selective to different reward types, we set out to test this prediction using data from Lak et al. (2014). In this experiment, a monkey was presented with 5 stimuli associated with different volumes of juice or different amounts of banana (Figure 3A). In the first part of the experiment, on each trial, the monkey was making a choice between two stimuli, and obtained corresponding rewards. Choices on these trials were used to estimate subjective utility of rewards associated with each stimulus. In the second part of the experiment, on each trial the monkey was presented with a single stimulus, and received corresponding reward. During these trials, the activity of dopamine neurons was recorded. Figure 3B displays the subjective utility of the 5 rewards estimated by Lak et al. (2014) from the first part of the experiment. The monkey exhibited a clear preference between different amounts of juice and banana rewards. The utility increased with the amount of reward, and the two most preferred rewards were the largest volume of juice and the largest quantity of banana.

**Figure 3:**
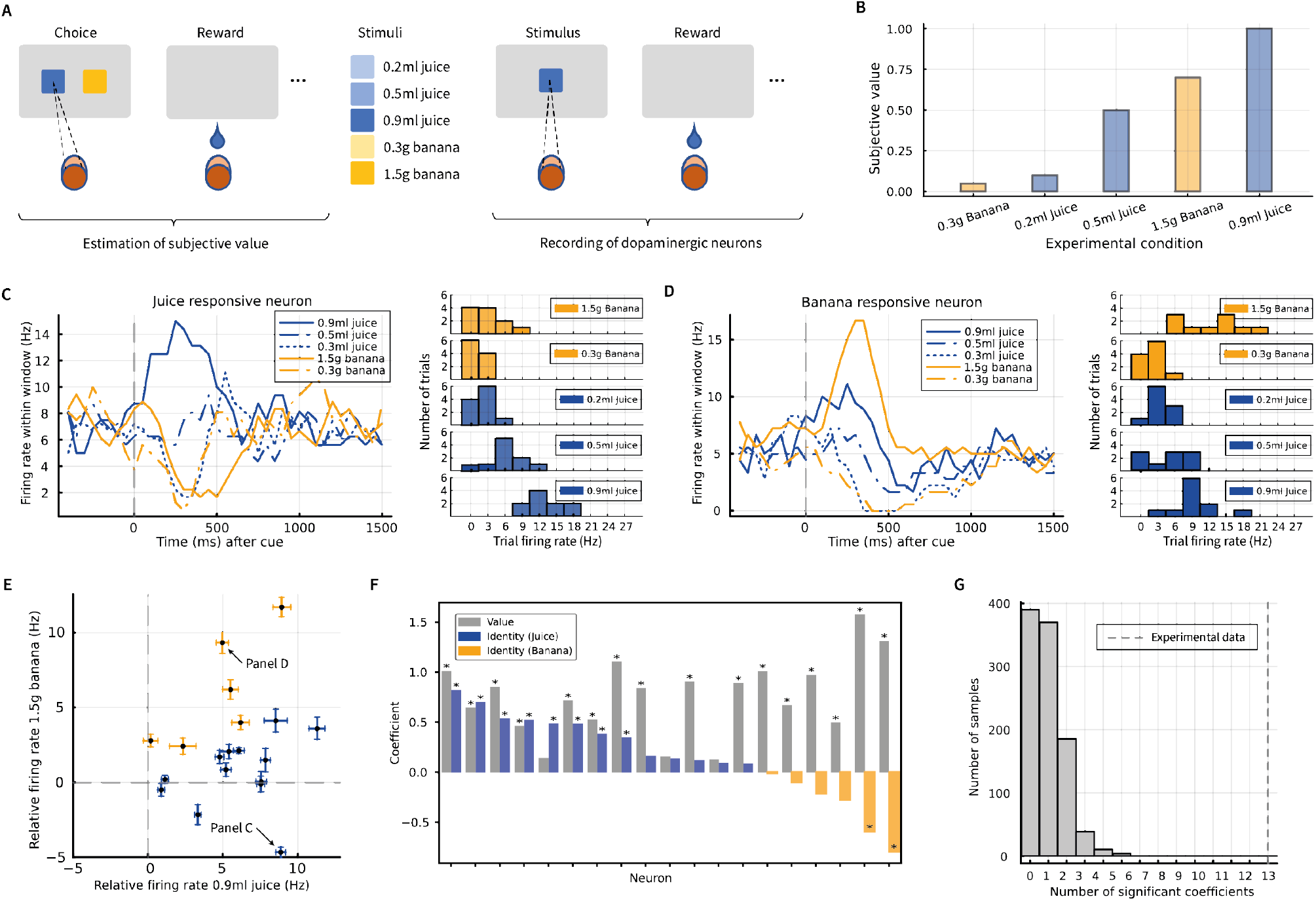
Results from analysis of single dopamine neurons in a monkey. **A**: The experimental paradigm used by (Lak et al., 2014). During recording, a monkey was presented with a visual stimulus indicating the reward type. We analyzed the responses of individual dopamine neurons in the period after the stimulus presentation. **B**: The subjective value determined by Lak et al. (2014) for each of the 5 conditions which could be presented to the monkey. **C, D**: Average neural firing rates during a trial for a highly juice-responsive neuron and a highly banana-responsive neuron. The left displays show the firing rate as function of time within a trial (see Methods), where time 0 corresponds to onset of the stimulus indicating which reward type will be presented. The right displays show histograms of firing rates within a window of 100-400 msec after stimulus onset for each condition for the corresponding neuron. **E**: Responses of individual neurons to stimuli predicting the largest food or drink rewards. Each dots corresponds to a single neuron, and its coordinates correspond to the average responses to either 0.9ml of juice (x-axis) or 1.5g of banana (y-axis). The response is computed as the firing rate within a window 100-400 after stimulus onset relative to the baseline 0-500ms prior to the stimulus onset. The error bars show the standard error of the mean, and are coloured according to neurons’ sign of identity regressor in panel F. **F**: The value and identity coefficients for a linear regression predicting the number of spikes in the post-stimulus period as a function of the subjective value and reward type. Stars indicate significant coefficients (*p <* 0.05 uncorrected, t-test for significance of regression co-efficient, two-sided). **G**: The number of neurons with significant identity coefficient when we repeated the analysis 1000 times and each time randomized the assignment of trials to conditions. Dashed line indicates the number of neurons with significant identity coefficient obtained in the analysis of the experimental data.

We re-analysed the activity of individual dopamine neurons, and found that indeed the majority of recorded neurons are differentially selective towards juice or banana, supporting the prediction of the Reward Bases model. Figure 3C shows an example of neuron whose activity is significantly modulated by the volume of juice predicted by the stimulus but little by the amount of banana. In particular, the firing rates differ substantially between stimuli predicting different volumes of juice, while they are similar for large and small amount of banana. Remarkably, the firing rate of this neuron decreases below baseline after cue predicting 1.5g of banana even though it was the second most favourite reward (Figure 3B). Such decrease is predicted by the Reward Bases model for a dopamine neuron encoding prediction error associated with drink, because banana provides less drink than the average across the rewards in the experiment. Figure 3D shows the activity of a neuron which is modulated by the amount of food more than by volume of juice. For this neuron, the responses to large and small amount of food are separated more than responses to different volumes of juice.

To visually illustrate the diversity of responses of dopamine neurons, Figure 3E shows the responses to the stimulus predicting largest amount of banana, against the stimulus predicting the largest volume of juice. If dopamine neurons encoded single reward prediction error signal from the standard TD model, then their responses to these two stimuli would be similar and positive, because these two rewards are preferred by the animal more than other outcomes, hence would produce positive prediction errors. By contrast, dopamine neurons produced diverse responses, and multiple neurons decreased their firing rates after the stimulus predicting 1.5g of banana.

To statistically test the selectivity of neurons for different reward dimensions, we quantified to what extent the activity of each dopamine neuron is dependent on subjective value and identity of reward. For each neuron, we fitted regression model predicting neuron’s firing rate after stimulus onset based on subjective value regressor and identify regressor (equal to 1 for juice and to −1 for banana). Figure 3F shows the estimated regression co-efficients for all neurons. Grey bars show the co-efficients for value, which are all positive, implying that the activity of all neurons is affected by the value. The blue bars show the co-efficients for identity that are positive, indicating the neurons responding more for juice, while yellow bars show the negative ones, indicating preference for banana. We found that 13 neurons possessed a statistically significant identity coefficients implying that their response depended on reward type. To investigate if such number of significant identify coefficients could arise by chance, Figure 3G plots the number of significant coefficients obtained in multiple repetitions of our analysis in which we shuffled assignments of trials to conditions. The figure shows that in all 1000 repetition the experimentally observed number of neurons with significant identity coefficient was smaller than in the experimental data, so the probability of obtaining 13 significant neurons under the null hypothesis is *p <* 0.001. This provides statistical support for model’s prediction that dopamine neurons are selective for different reward dimensions.

The above analysis of experimental data provides a partial support for the fundamental Reward Bases model introduced so far. As predicted, different dopamine neurons are selective for different reward dimensions. However, the fundamental version of the model assumes that each neuron should encode a single reward basis, while in the neural data this distinction is less absolute. Instead, neurons appear to be more or less selective for a reward type, but not entirely unselective for the other. Nevertheless, the model can be refined to capture such variation.

We consider a mixed selectivity model (details in Methods section) in which each dopamine neuron *k* computes a mixture or a weighted sum of prediction errors associated different reward bases 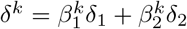. Each dopamine neuron in the model has different weights 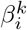 for different reward bases. Such computation could arise from connectivity pattern shown in Figure 4A, where each dopamine neuron receives inputs from reward and striatal neurons associated with distinct reward bases with particular weights. The value bases are then learned in this model through the standard TD error rule using weighted sums of the prediction errors. We simulated a model that contained the same number of dopamine neurons, as in the data in Figure 3, and encoded the mixtures of prediction errors with weights estimated from individual in biological neurons. To estimate the weights, for each dopamine neurons we fitted a regression model that predicted firing rate after the stimulus onset based on prediction error for the two reward bases. The obtained regression co-efficient are plotted in Figure 4B, providing another illustration for the diversity in selectivity of dopamine neurons. The estimated regression coefficients were used as weights of prediction errors 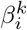 of the corresponding neurons in the model. Figure 4C shows that the resulting model can learn accurate value functions in a simulated version of the task of Lak et al. (2014).

**Figure 4:**
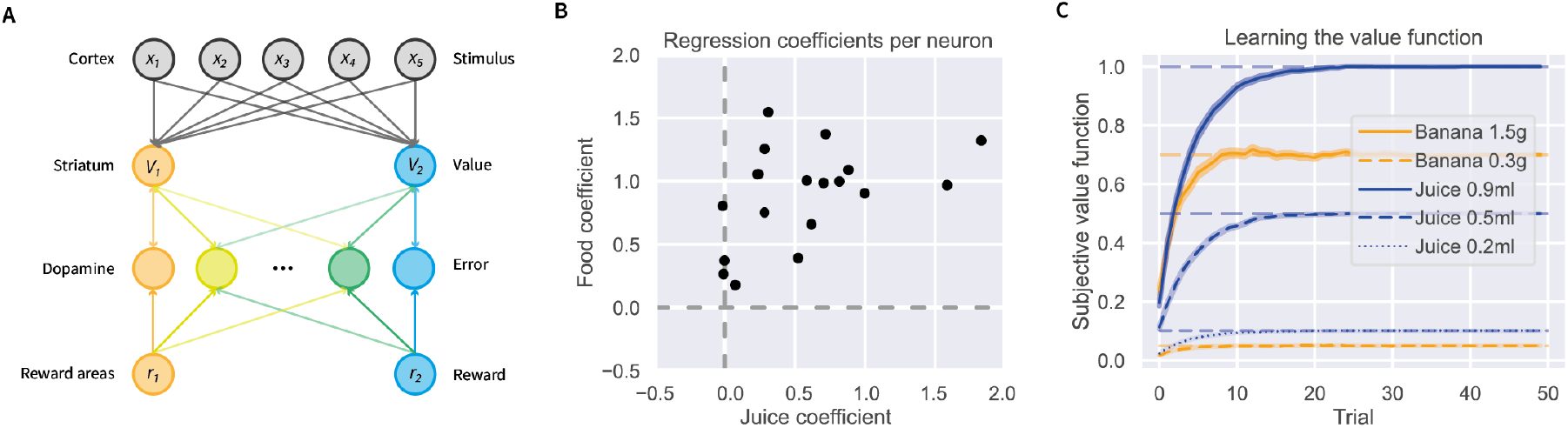
Mixed selectivity model. **A**: A possible neural circuit giving rise to Reward Bases model with dopamine neurons selective for a mixture of prediction errors. Since we do not have data from striatal neurons, for simplicity we consider an idealized model in which different reward dimensions are encoded by separate striatal populations, and two such populations are shown in orange and blue. Thickness of lines corresponds to the strengths of the connections, and the gradation of colour of dopamine neurons indicates their degree of selectivity for the two reward bases. **B**: Regression co-efficients indicating to what extend individual dopamine neurons encore prediction errors associated with the two reward bases. Each dot corresponds to one neuron. Its co-ordinates are the regression co-efficients estimated from biological neurons, and are used as weights for different prediction errors encoded by neurons in the model. **C**: Mean values of stimuli learned in a simulated version of the task of Lak et al. (2014). The shaded areas represent the standard error of the mean. We only plotted the meaningful (non-zero) learnt values since the values of the banana reward condition under the juice reward basis were 0 and vice-versa. Horizontal lines show the true subjective values (also shown in Figure 3B), indicating that the mixed selectivity model can reliably learn the correct values for each condition.

Simulations in Figure 4C establish that the model with mixed selectivity of dopamine neurons can learn value bases similarly as the original Reward Bases model. Since we do not compare the model to data from individual neuron in the remainder of the paper, we will not include mixed selectivity in the remaining simulations for simplicity.

### Dopaminergic activity reflects instant revaluation

In this and the next sections we demonstrate that the Reward Bases model captures instant revaluation after changes in physiological state seen in dopaminergic activity and behaviour, which cannot be described by the standard TD-learning. Here, we focus on qualitatively matching the dopaminergic responses in a reward devaluation task in which rats could press levers associated with either a food or a sucrose reward under varying conditions of selective satiation (Papageorgiou et al., 2016). The experimental data we modeled were taken from forced trials in which only one of the levers delivered rewards. A schematic of the experimental paradigm of Papageorgiou et al. (2016) can be seen in Figure 5A. At the onset of each forced trial, animals were presented with a cue indicating which of the two levers can be pressed to obtain a reward. Each lever typically delivered a particular type of reward, but on some trials it could give either four times the amount of reward (MORE condition), or the other reward type (SWITCH) condition. In the devaluation condition, the animals were fed to satiation in one of the reward types but deprived of the other. Fast-scan cyclic voltammetry was used to measure dopamine levels in the nucleus accumbens core, part of the ventral striatum.

**Figure 5:**
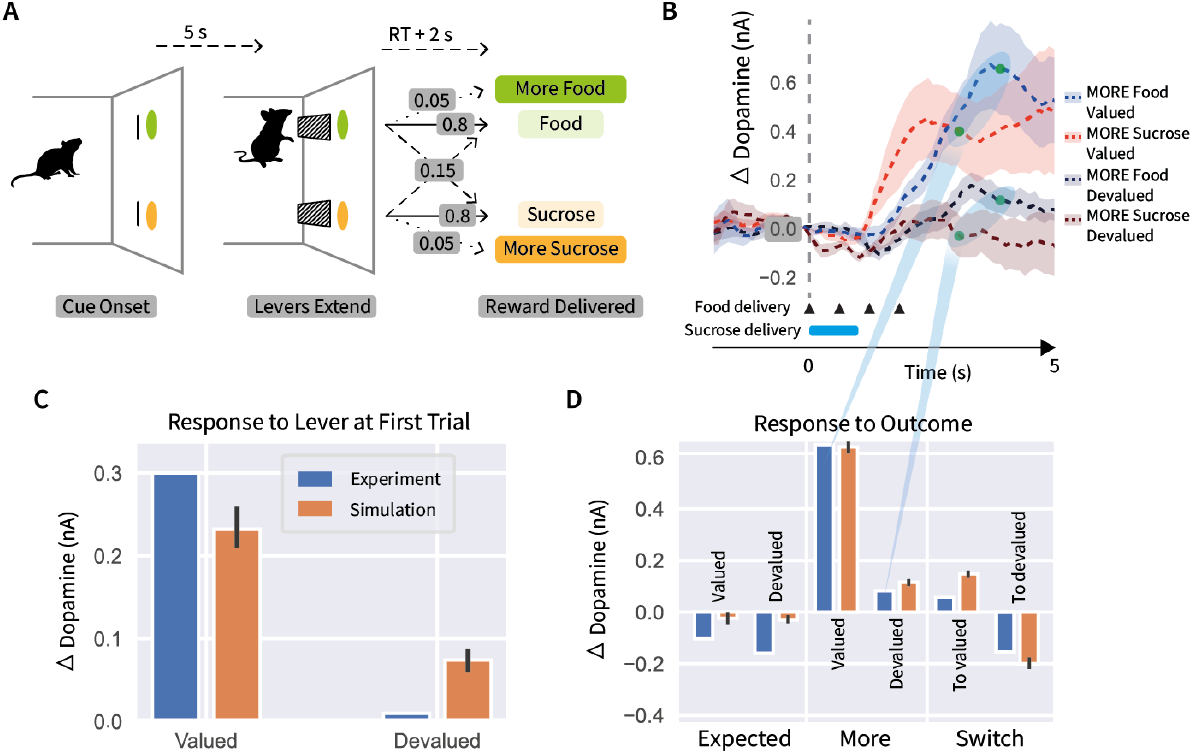
Dependence of dopaminergic responses on physiological state. **A**: Sequence of events during a trial in the experimental paradigm of Papageorgiou et al. (2016). At the onset of each trial, animals were presented with a cue indicating which of the levers trigger reward delivery. After a delay of 5s lever(s) extend, and following Reaction Time (RT, i.e. time to press a lever) and further 2s a reward is delivered. A particular type and size of the reward was delivered with a probability indicated by labels next to arrows. **B**: Dopaminergic activity in the MORE condition (re-plotted from Figure 2c in the paper by Papageorgiou et al. (2016)). In this condition the reward delivery was extended in time: 4 pieces of food were delivered every 600ms as indicated by triangles, while sucrose was delivered for 1s as indicated by a blue bar. Model simulations were compared with dopamine levels 2s after the end of reward delivery (indicated by green dots), which were averaged across food and sucrose to give “experimental data” in panel D, as illustrated by the blue shading to panel D. **C**: Comparison of simulated dopamine level against data after extension of lever indicating which reward are likely to be available, taken on the *first trial* just prior to reward delivery. The experimental data corresponds to dopamine level 2s after lever extension (data was read out from dashed curves in Figure 4a in the paper by Papageorgiou et al. (2016)). **D**: Comparison of simulated results against dopaminergic responses to reward delivery. The experimental data in EXPECTED and SWITCH conditions corresponds to dopamine level 2s after reward delivery (read out from Figures 2d-e in the paper by Papageorgiou et al. (2016)).

To capture dopaminergic activity when animals are deprived of a particular resource (Cone et al., 2016; Papageorgiou et al., 2016), models need to assume that dopaminergic signals encoding prediction errors are modulated by animal’s physiological state (van Swieten & Bogacz, 2020). For example, Figure 5B shows responses to different rewards in MORE conditions in different states in the task illustrated in Figure 5A. The dopamine level increases after receiving more of food (or drink) than expected only if the animal was hungry (or thirsty), suggesting that the dopamine neurons encode prediction errors scaled by the corresponding physiological need. To account for this data, a modification needs to be made to Reward Bases model: it needs to be assumed that dopamine neurons associated with reward basis *i* encode 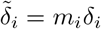. In the Methods we further motivate and describe such model with state-dependent prediction error, in which 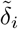 also drive synaptic plasticity. Note that this modification does not affect predictions tested or simulations earlier in the manuscript, because in the experiment of Lak et al. (2014) the animals were motivated to acquire both food and fluid (so 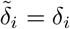 for *m*_*i*_ = 1), and in the simulation in Figure 1 the learning took place while *m*_*i*_ = 1.

To model these data, we simulated an agent interacting with the Papageorgiou et al. (2016) paradigm illustrated in Figure 5A. The agent was trained with two reward bases, one which gave out 1 reward for each food reward and 0 for sucrose, and a sucrose basis with the opposite reward schedule. To simulate testing in devaluation sessions where the animals were fed to satiety in one reward type but deprived of the other, the reward basis weights *m*_*i*_ of the agent were fitted to the data to determine how the reward weights changed during devaluation. Since voltammetry measures relative changes in extrasynaptic dopamine, we use our model to make predictions of total dopamine release, which we take to be the sum of the dopamine neurons, i.e, 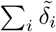, summing across the contributions of all reward bases.

A key result of Papageorgiou et al. (2016) is shown by blue bars in Figure 5C, visualizing the dopamine response to the lever extension on the *first trial* after devaluation, before the animals have any experience with the devalued reward. The response to a lever associated with the devalued reward is lower than for the valued option even on the first trial (Papageorgiou et al., 2016). This finding cannot be accounted for by standard TD learning, because the prediction errors following a cue reflect the values of the cues, which in TD-learning model are only updated following reinforcements, but since devaluation the animals did not receive any rewards. Reward Bases model, however, straightforwardly predicts these results. This is because the weighting coefficient *m*_*i*_ is lower for the devalued reward, thus the prediction error 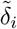 associated with devalued option is also reduced, even before the devalued outcome is delivered.

Figure 5D shows that the Reward Bases model can capture key qualitative pattern seen in dopaminergic responses to outcome. When the most common reward type is delivered the observed dopaminergic responses slightly decreased, and in the model the sum of prediction errors is slightly negative, because MORE or SWITCH trial did not occur. In the MORE condition, the dopamine response is much higher when the large amount of valued reward is delivered, and the model reproduces this pattern because the positive prediction error 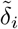 is scaled by a larger weight for the valued reward. In the SWITCH condition, the dopaminergic response is higher when valued rather than devalued reward is given, and the model reproduces this pattern because the positive prediction error caused by switch to valued dimension is scaled by a larger weight.

### Behaviour reflects instant revaluation in novel physiological state

In this section we show that Reward Bases model can account for instantaneous changes in behaviour following a novel physiological state, which cannot be described by the TD learning. As mentioned in the Introduction, classic experiment that demonstrated instant reward revaluation abilities was conducted by Robinson and Berridge (2013). They utilized a Pavlovian association paradigm in which rats were repeatedly exposed to one of two cues – associated with either intra-oral delivery of sucrose (pleasing) or salt (aversive) solution. The rats were trained in a food-restricted state but when injected with aldosterone and furosemide which mimics severe salt deprivation, immediately responded positively to the salt cues (Figure 6A). These results imply that the rats clearly possess the ability to perform instant generalization to update associations upon physiological change with no direct experience of the positive rewards associated with the change. This phenomenon cannot be explained by the TD model, since the animals never experience the salt solution as rewarding, and have no opportunity to update their value function. These results can, however, be directly explained by the Reward Bases model.

**Figure 6:**
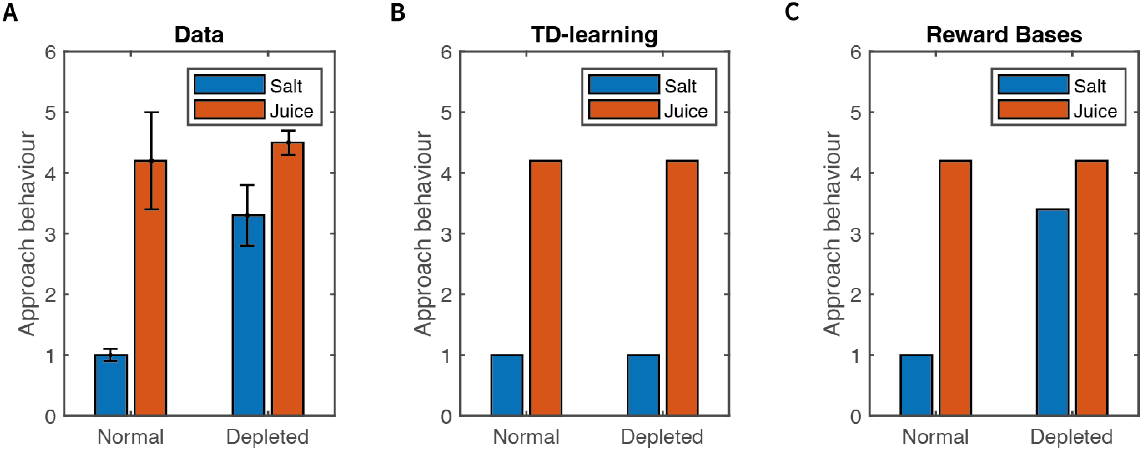
Effects of salt deprivation on behaviour in the study of Robinson and Berridge (2013). **A**: Experimental data showing approaches or appetitive actions towards the salt and juice lever in the homeostatic (normal) and salt-deprived conditions (data is replotted from Figure 3C in the study of Robinson and Berridge (2013).). **B**: Simulation of TD-learning. **C**: Simulation of Reward Bases model.

Since salt is a physiologically important quantity (Krause & Sakai, 2007), we assume the animals would maintain a ‘hardwired’ reward basis computing a salt value function during the Pavlovian association phase. However, upon the injection of the aldosterone and furosemide, the Reward Bases model can dynamically modulate its (negative) salt value function with its physiological state to perform instant revaluation of its associations with the salt lever. This can occur even in the absence of any positive reward signal obtained by experiencing the salt, since the weighting coefficients *m*_*i*_ act directly on the value bases, and thus the value bases themselves do not have to be updated. We demonstrate this instant generalization capability of the Reward Bases model by replicating the paradigm of Robinson and Berridge (2013) in simulation and matching the experimental behavioural results obtained.

In simulation, the rats were exposed to interactions with the salt or juice levers (at random) and learnt a value function of being near the salt or juice levers. We assumed a linear proportional relationship between the learnt value function and degree of appetitive behaviour towards the lever. Figure 6B-C shows that in the homeostatic (normal) condition, both TD learning and Reward Bases model are able to reproduce the behaviour of the animal, while in the sodium depletion condition, the Reward Bases agent can instantly generalize to match the behaviour of the agent, thanks to the weighting coefficients *m*_*i*_ while the TD agent has no mechanism to vary its value function according to its physiological state, and so simply predicts that the salt lever will remain aversive to the animal in the sodium depleted condition.

### Comparison with alternative models

In this section we perform a computational evaluation of our model against alternative models in a room navigation task introduced in Figure 1. We demonstrate that the Reward Bases model performs equivalently to the successor representation (Dayan, 1993) at a smaller computational cost, while it outperforms TD learning and the homeostatic model (Zhang, Berridge, Tindell, Smith, & Aldridge, 2009).

Our test environment is a 6 × 6 grid-world room (Figure 1). The goal is, when started in a random position in the room, to reach a specific type of object as fast as possible. There were three objects of different types that were positioned randomly at the start of simulation. Agent training was separated into episodes such that whenever the agent reached the valuable object, the episode would end, and the agent would restart in a randomly chosen square of the room. At each moment of time only one object was valuable, and after a certain number of trials objects’ desirability was reversed – making one of the three objects valuable while demoting the others.

In Figure 7A, we plot the performance of the models in sample simulation. As expected, the Reward Bases agent performs better that TD-learning which needs to completely re-learn value function after each reversal. We also simulated the ‘homeostatic model’ of Zhang et al. (2009), which modulates the expected instantaneous rewards on the next state according to physiological state (see Methods for details) and which has been proposed as a model of the instant reward revaluation observed in animals. Since this model directly modulates predicted rewards, it can solve single-step tasks such as the Dead Sea-salt experiment but cannot generalize behaviour across multi-step tasks since it does not modulate *value functions* which is necessary to correctly adapt long-term behaviour upon changing the reward function.

**Figure 7:**
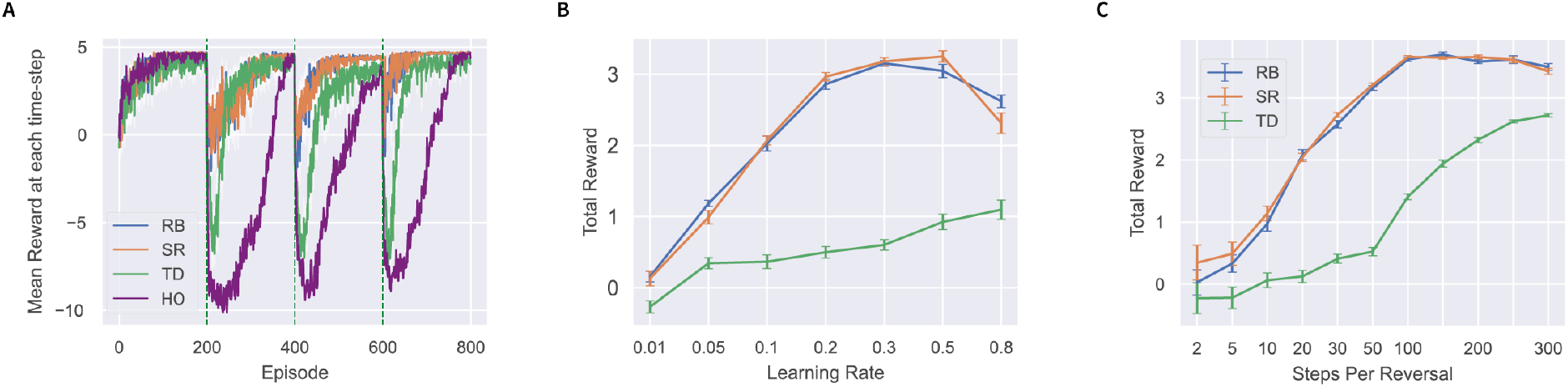
Comparison of model performance in a room navigation task. **A**: Comparison of the reward obtained by the Temporal Difference (TD), the Reward Bases (RB), the Successor Representation (SR), and the Homeostatic (HO) agents on an example run of the room task. The vertical dashed lines represent the reversals. **B**: Performance obtained by RB, TD, and SR agents over a range of learning rates with 50 episodes between each reversal. **C**: Performance obtained by RB, TD, and SR agents over a range of trials between reversal. A learning rate of 0.1 was used. All error bars represent standard deviation over 10 runs.

Additionally, we compare our model to successor representations which can generalize correctly across changing reward functions. Successor representations (Dayan, 1993) is an alternative model-free RL method that also enables instantaneous generalization across changing reward functions. They work by learning a ‘successor matrix’ based on discounted state occupancies (see Methods section for mathematical details) which can then be combined with the reward function to yield the value function. Figure 7A shows that the successor representation has very similar performance as Reward Bases. Both agents rapidly adapt to reversals but still also require some retraining after reversals likely due to the approximate nature of the learnt value functions or successor matrix given their limited number interactions with the environment. In performance on this task, the Reward Bases and successor representation agents both achieve approximately equal performances across a wide range of parameter settings (Figure 7B-C). In Methods we show mathematically that the Reward Bases model is closely related to the successor representation and, indeed, can be intuitively thought of as a compressed successor representation tuned only to relevant reward dimensions. Therefore, the Reward Bases and successor representations have approximately equivalent capabilities, but Reward Bases model stores much less information. The value bases store *X × N* numbers, where *X* is a number of states, while *N* is the number of reward bases, while the successor matrix stores *X* × *X* numbers. This difference is already substantial for the small task presented here, where with three reward bases, the number of values stored by the Reward Bases algorithm was 36 × 3 = 108 while the successor matrix required 36 × 36 = 1296. This advantage will generally only increase with state-size, because the memory cost of the Reward Bases model scales linearly with the state-size, while for the successor representation it scales quadratically.

## Discussion

In this paper, we have proposed a novel mechanism for instant generalization to changing reward functions in RL tasks which relies solely on model-free TD learning methods. Crucially, our results demonstrate that behavioural flexibility in the face of changing reward functions does not require model-based methods as argued by Dayan and Berridge (2014) but instead can be handled in a purely model-free manner using a straightforward extension to the classical TD learning method which is already widely believed to be implemented in the basal ganglia. Moreover, this method is extremely straightforward and can be applied to any RL model which estimates values of states or actions.

We have also verified the key experimental prediction arising from our model and demonstrated that different dopamine neurons are selectively sensitive to different reward types. This observation is consistent with a recent study showing that separate groups of dopamine neurons increased their tonic firing after intragastric infusion of either food or water (Grove et al., 2022). That study also found no correlation across dopamine neurons between responses to food and water, which parallels the lack of correlation in Figure 3E. Our analysis extend the results of Grove et al. (2022) by showing that different dopaminegic neurons also produce phasic responses after stimuli predicting different types of rewards, and the these responses scale with the amount of a particular reward type. Our results are also consistent with a recent observation that activity of dopamine neurons depends on separately computed predictions for different types of rewards (Takahashi et al., 2023). Our analysis additionally shows that if the rewards are more distinct, then dopamine neurons themselves are selective for reward type. Furthermore, our results are consistent with a study of Huang and Grabenhorst (2023) showing that primate behaviour is well described by a model also including prediction errors for different reward dimensions. The model of Huang and Grabenhorst (2023) learns instantaneous expected reward in multiple dimensions, and our paper additionally shows how such model can be extended to learn discounted future reward, and how such learned values may be combined to enable flexible behaviour.

Our results add support to a general theory that the striatum is parcellated into modules predicting different quantities, and the dopamine neurons projecting to a given parcel of the striatum compute the error in prediction made by this parcel (Bogacz, 2020). In the case of Reward Bases model, different parcels located in ventral striatum compute different value bases, but other striatal regions may compute different quantities, and consequently corresponding dopamine neurons may encode errors in different predictions. For example, dopamine neurons in the tail of the striatum were proposed to encode prediction errors specifically for aversive stimuli (Brischoux, Chakraborty, Brierley, & Ungless, 2009; Watabe-Uchida & Uchida, 2018), or action prediction errors (Greenstreet et al., 2022). Moreover, dopamine neurons have been shown to encode errors in predictions of sensory features of reward in addition to value (Takahashi et al., 2017). There is also growing evidence that dopamine neurons are sensitive to various aspects of the state-space such as locomotion and kinematic behaviour (Barter et al., 2015; Dodson et al., 2016; Engelhard et al., 2019; Kremer, Flakowski, Rohner, & Lüscher, 2020) as well as choice behaviour (Coddington & Dudman, 2018; Parker et al., 2016) and indeed that this heterogeneity is topographically organized due to spatially organized cortical projections (Engelhard et al., 2019; Howe & Dombeck, 2016). Additionally, recent theoretic work has demonstrated that RL agents including separate modules predicting reward in different dimensions have advantages in task requiring acquisition of multiple resource types (Dulberg, Dubey, Berwian, & Cohen, 2022).

The existence of striato-dopaminergic modules seems to contradict a common belief that activity of dopamine neurons is transmitted throughout the whole striatum. This belief originates from the observation that dopamine receptors are located outside synapses so they can be only activated by dopamine diffusing through the space (Liu, Goel, & Kaeser, 2021). However, Liu et al. (2021) point out that only a small volume of space around the axonal release site (within 1*μm*) is likely to reach sufficient dopamine concentration for receptor activation. So although dopamine release is not targeted at single synapses, it has substantial spatial precision. Moreover, despite dopamine neurons being known to have exceptionally large axons, Matsuda et al. (2009) reported that axonal bush of a single dopamine neuron covers on average 2.7% of volume of striatum in the corresponding hemisphere, and this area in concentrated to a particular region of striatum. Hence the existence of modules within the striatum modulated by distinct sets of dopamine neurons is anatomically plausible.

It is useful to clarify the distinction between the Reward Bases model and distributional RL (Dabney et al., 2020). That model assumes that the value function is represented by the probability distribution of expected reward. In that model separate parcels learn different percentiles of reward distribution and the heterogeneity of dopamine neurons arises as they encode prediction error associated with different percentiles of the distribution. By contrast, the Reward Bases model proposes heterogeneity in dopamine neurons in another orthogonal dimension – that of reward type. It is also possible to straightforwardly combine distributional RL and the Reward Bases models by estimating a distributional value estimate for each value basis in parallel.

Given the importance of being able to appropriately select actions according to physiological state, similar mechanism to those considered in this paper may also operate in simpler animals that do not have the basal ganglia. In particular, there is a strong match between the neuroanatomy postulated by the Reward Bases model and that found in the mushroom body of fruitflies (drosophilia). This region includes Kenyon cells encoding sensory information, and Mushroom Body Output Neurons (MBON) which are mutually connected with dopamine Neurons (DAN) (Li et al., 2020). A large body of evidence, strongly suggests that the MBONs and DANs are specialized and separated into different zones or compartments which respond to and represent rewarding aspects of specific stimuli, and connectivity between MBONs and DANs is mostly within a given compartment (Aso et al., 2014; Li et al., 2020; Otto et al., 2020; Owald & Waddell, 2015; Vogt et al., 2014). Moreover, experimentally it has been shown that specific DANs respond to specific reinforcement types such as sugar (Burke et al., 2012; Huetteroth et al., 2015), water (Lin et al., 2014; Senapati et al., 2019), courtship (Cheriyamkunnel et al., 2021; Keleman et al., 2012) and aversive stimuli (Aso et al., 2012; Owald & Waddell, 2015; Perisse et al., 2016; Riemensperger, Völler, Stock, Buchner, & Fiala, 2005), and appear to be instrumental in learning associations based on these specific reinforcement type. Hence if we associate sensory state *x* with Kenyon cells, value bases *V*_*i*_ with the MBONs and the prediction errors *δ*_*i*_ with the DANs, then the circuitry in the mushroom body appears to almost precisely fit a direct implementation of the Reward Bases model.

In this paper, we have simply assumed that the weighting coefficients *m*_*i*_ are known to the agent, but the brain would have to determine the correct weightings, and transmits these values to the dopaminergic system. One way to bring an organism towards a homeostatic set-point is to set the weights *m*_*i*_ for a specific homeostatic variable proportional to the deviation of that variable from an optimal set-point (van Swieten & Bogacz, 2020). Such a model is closely related to drive reduction theory (Hull, 1943; Juechems & Summerfield, 2019; Keramati & Gutkin, 2011) which fits closely with notions of homeostasis and allostasis in biology.

The present study provided initial evidence for learning value in multiple reward dimensions in the basal ganglia. Future studies are needed to identify which dimensions of reward are represented by the dopamine neurons, whether neurons encoding specific dimensions are topographically organized, and if dopamine neurons can learn to represent novel reward dimensions.

## Methods

### Temporal Difference learning

In multiple simulations we included the TD learning model for comparison with the Reward Bases model. In the TD model, the value function is learned by directly updating the estimated value of the current state (Sutton & Barto, 2018),

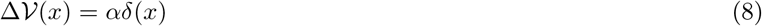

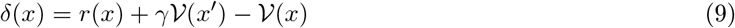

This learning rule is analogous to that in the Reward Bases model, but is based on the total reward *r*(*x*) rather than reward bases.

### Analysis of activity of individual dopamine neurons

For a full description of the paradigm and data acquisition, please refer to the original paper (Lak et al., 2014). The data from Lak et al. (2014) consists of series of spike-times for each neuron for each trial and the associated condition. There were five conditions corresponding to the monkey being presented with 1.5g banana, 0.3g banana, 0.9ml juice, 0.5ml juice, and 0.2ml juice. We removed trials with less than 5 total spikes over the 10 second trial due to a lack of signal. The average number of spikes per trial was 48.7 with a standard deviation of 14.6 so *<* 5 spikes was more than 3 standard deviations from the mean. In total only 7 out of 1225 trials were removed. We also aligned all spike-times so that all trials started with the stimulus presentation at 0ms. Besides this, we performed no other data-preprocessing.

To obtain the subjective values plotted in Figure 3B, we read off the values from the subjective value plot of Figure 4 in Lak et al. (2014). We then shifted the obtained values by +0.5 so as to move the range of subjective values to lie between 0 and 1. This has no impact on the relative ranking or differences between conditions but makes the resulting regression coefficients more interpretable.

In Figures 3C and D, we plot the average firing rate over the trials of each condition for a highly selective ‘food’ or ‘juice’ neuron. To obtain a smoothed number of spikes, we counted the number of spikes within a set of overlapping windows, each 100 msec long and starting every 50 msec. To convert from the raw number of spikes in the window to an average firing rate, we simply divided the number of spikes by the window size (in seconds). Since the window size was 0.1 seconds (100 msec), this was equivalent to multiplying by 10. We plot the average firing rate of the neuron in each condition. We aligned the timestamps of each trial so that the condition cue was presented at time 0. In the accompanying histograms, we show the distribution across trials of neural firing rates within a window of 100-400 msec after stimulus onset for each condition. The firing rates are obtained as before by dividing the number of spikes within the window by the window size to obtain an extrapolated firing rate.

To statistically quantify if individual dopamine neurons are selective for a particular reward type, we employed a regression analysis. For each neuron we fitted a model predicting the number of spikes emitted in each trial within a window of 100-400 msec after stimulus onset, based on value and identity regressors. The value regressor on a given trial was assigned to the subjective value of reward on that trial found by Lak et al. (2014) and plotted in Figure 3B. The identity regressor was coded such that +1 indicated a juice trial and −1 indicated a banana trial.

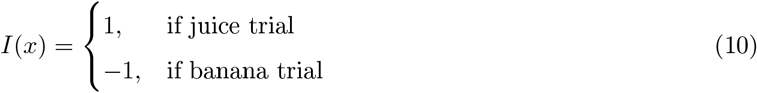

Mathematically, we can formalize the linear model we tested as

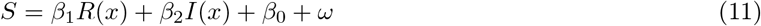

where *S* is the number of spikes within the window, *R*(*x*) is the value regressor, *I*(*x*) and is the identity regressor. *β*_1_ and *β*_2_ are the regression coefficients, *β*_0_ is the intercept term and *ω* is the noise term in the linear model. Before performing the regression, we normalized the regressors such that they had a mean of 0 and a standard deviation of 1. This did not affect the significance of the regressors but increased the interpretability of the magnitude of regression coefficients. Figure 3F plots the regression coefficients *β*_1_ and *β*_2_ for each neuron. To validate the statistical significance of our results of 13 significant identity coefficients, we investigated empirically how many significant coefficients we should expect with a random assignment of conditions to trials. We ran 1000 independent regressions identical to that described above, except where we randomized the assignment of conditions (and hence regressor values) to trials. For each of our trial regressors, each trial was assigned a condition by sampling from the probability distribution of the fraction of trials in each condition in the true dataset. The conditions were sampled with replacement. The number of neurons with significant identity regressor across 1000 repetitions are plotted in Figure 3G.

### Mixed Selectivity Model

We constructed a model in which each dopamine neuron encoded a weighted sum of prediction errors with weights estimated from individual biological neurons. These weights were estimated by fitting a regression model predicting the total number of spikes *S* in a window of 100-400msec after stimulus onset, based on a juice subjective value and a food subjective value regressors. Specifically, we implemented the following linear model,

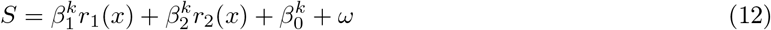

where *r*_1_ and *r*_2_ are the reward bases for the banana and juice regressors, 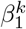 and 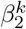 are the regression coefficients of dopamine neuron *k* for each reward basis while 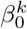 is the intercept term and *ω* represents the noise term. The juice reward basis here assigned the subjective values associated with the juice for the juice conditions and 0 for the food conditions. Conversely, the banana reward basis here assigns the subjective values associated with the banana for the banana conditions and 0 otherwise. For instance, we can formally define the juice reward basis *r*_1_ for condition *x* as,

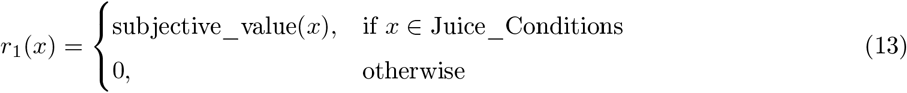

The subjective values are those taken from (Lak et al., 2014) and replotted in Figure 3B. Note that if we used regressors equal to prediction errors rather than *r*_*i*_, the resulting coefficients 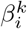 would be identical, because the prediction errors just differ from *r*_*i*_ by a constant (expected reward) and these constants are incorporated into 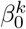 in our regression. To enable interpretation of the resulting coefficients, the regressors were then normalized to have mean 0 and a variance of 1. The resulting regression coefficients are plotted in Figure 4B.

We simulated a model with *K* dopamine neurons which each encoded separate reward prediction errors *δ*^*k*^(*x*). An abstract version of the experimental paradigm of Lak et al. (2014) was simulated where at each trial one of the 5 stimuli was presented. The goal of the experiment was to learn the values of each stimulus. The stimuli were randomly selected across 100 trials. In our simulation, we only considered two reward bases, one pertaining to banana rewards (food) and one to juice rewards (water). Whenever a stimulus was presented, its subjective value (as plotted in Figure 3B) was provided to all dopamine neurons *k* = [1 … *K*], weighted by coefficients 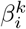 describing to what extent neuron *k* is selective for reward dimension *i* (equal to the regression coefficients for each neuron shown in Figure 4B). Each dopamine neuron also received estimated value and calculated a prediction error:

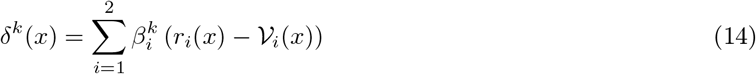

This prediction error term contains no future value term *γV*_*i*_(*x*^*′*^) since our task lasted for just a single step. Then, the value bases were updated according to a weighted sum of the dopaminergic responses of each neuron

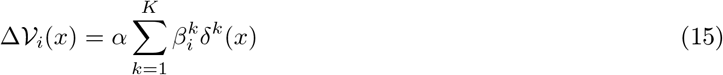

where *α* is a scalar learning rate which was set to 0.1. To understand why such model can learn the correct value bases, please note that it implements a gradient descent on summed square prediction error of all dopamine neurons:

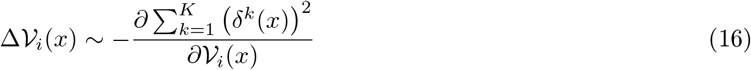

The simulation was repeated 1000 times, and resulting mean values are shown in Figure 4C.

### Model with state-modulated prediction errors

In the model described at the start of the Results section, the reward bases *r*_*i*_ and the weighting coefficients *m*_*i*_ are decoupled. The rewards and reward prediction errors are still computed and the value basis is still updated even if that basis is not valued at the current moment. This is mathematically optimal given no computational or resource constraints since it enables flexible and instant generalizations to the greatest possible range of reward revaluations. However, there is a growing body of research that suggests that this is not the case in the brain since animals only appear to learn value functions of tasks when they are important to the animal and thus have motivational salience. For instance Aw, Holbrook, de Perera, and Kacelnik (2009) show that when fish are first exposed to a stimulus when hungry, they will chose it substantially more often over an equally palatable alternative than when they were exposed to the stimulus when sated. This implies that there is a direct modulation of the values learnt during training based on the current physiological state which is maintained even after the physiological state has changed. Cone et al. (2016) has similarly shown that physiological state can modulate dopaminergic prediction error responses such that when the animals are trained to receive food rewards after a cue in a ‘depleted state’ i.e. they are hungry – in testing, dopamine neurons respond to the cue (CS) and not the food reward (US) as in the classic results of Schultz et al. (1997). However, when the animals are trained in a sated state, when tested the dopamine neurons respond only to the food reward (US), ‘as if’ they had not learned the task at all.

The version of Reward Bases model described at the start of Results cannot explain these findings since it decouples dopaminergic activity – and hence value function learning – from the physiological state such that the state is only used to dynamically modulate the weighting of the value function bases during action evaluation. However, as shown by van Swieten and Bogacz (2020) such effects can be straightforwardly explained by redefining the dopaminergic prediction errors such that they are also modulated by the physiological state. Effectively, this means that the ‘learning rate’ of the TD update is modulated by the physiological state variable such that animals learn much more rapidly when they are ‘depleted’ and much less rapidly when they are sated. van Swieten and Bogacz (2020) demonstrated that such state-modulated prediction error can also be viewed as error in prediction of subjective utility of rewards given specific mathematical assumptions about the form of the utility function.

These insights can be straightforwardly incorporated into the Reward Bases model by adding motivational salience to the prediction errors directly over and above the modulation of the value bases, which greatly aids its neural plausibility. Mathematically, this model can be expressed by defining a ‘modulated prediction error’ 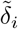,

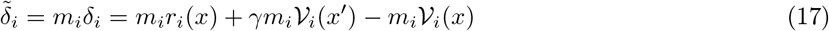

The TD update can then be derived as a gradient descent on the squared prediction errors,

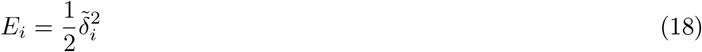

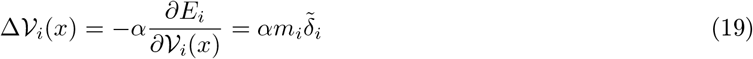

This results in a modified TD learning rule for the value bases which effectively defines an adaptive learning rate schedule where the learning rate depends on the physiological state. Although slightly complicating the algorithm, this approach has potential advantages for the brain. It provides a natural scaling of the learning rate with physiological stress, so that the learning rate is higher when the physiological state is more perturbed and hence the *m*_*i*_ weightings are higher. This has clear advantages since it is important to learn fast in such cases. On the other hand, having a reduced (or no) learning rate in the case of satiation may also be beneficial. Reducing the learning rate may reduce the metabolic cost of making the updates, since less synaptic plasticity is required and the brain has been heavily optimized by evolution to minimize energy expenditure (Sterling & Laughlin, 2015).

### Modelling the dependence of dopaminergic response on physiological state

The Reward Bases agent was trained in a simulated version of the task paradigm used by Papageorgiou et al. (2016) and graphically described in Figure 5A. The agent maintained two reward and value bases 𝒱 _*i*_(*x*) over the two states of the experiment *x*_*k*_ corresponding to pressing the two levers. On each trial the agent was choosing between two options, with probability of selecting option *k* equal to:

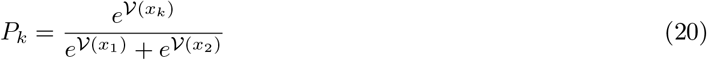

In the above equation, 𝒱 (*x*_*k*_) was computed from Equation 5 with *m*_*i*_ = 1. After choice, there was a 80% chance of getting corresponding reward type, a 15% percent chance of switching to the other reward type (SWITCH condition) and a 5% percent chance of getting 4 outcome units (MORE condition). Consequently both value bases for chosen option *k* were modified according to

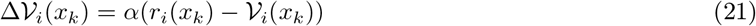

In the above equation, the discounted future reward was not included, because no further reward was expected, and learning rate was set to *α* = 0.1. Simulation of learning were repeated 3 times with unique seeds of random number generator, and within each repetition the value functions were learned over 500 trials.

The learned value bases were used to simulate dopaminergic responses in all experimental conditions in Figure 5C and D. Due to similarity of dopaminergic responses to food and water when these reward were valued (or devalued) seen in Figure 5B, we summarized these data by the dopamine concentration averaged across valued (or devalued) trials, as illustrated by blue shading from Figure 5B to Figure 5D. The dopaminergic responses were simulated as the sum of the weighted prediction errors, in accordance with the modulated prediction error model presented in the previous section. That is, we identify dopaminergic response in state *x* as

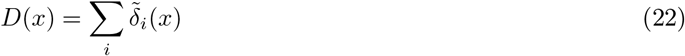

Without the loss of generality, we assumed that food was a valued option, while sucrose was devalued. Thus for example, the response to valued lever (Figure 5C) was computed by substituting Equation 17 into Equation 22

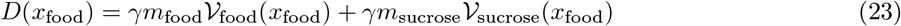

where *x*_food_ denotes the lever commonly associated with food. For simplicity we set *γ* = 1 in the simulations. The response to devalued lever *D*(*x*_sucrose_) was computed analogously. Similarly, the response to the expected valued reward (Figure 5D) was computed as

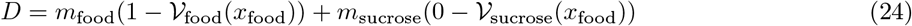

Responses to other outcomes were computed analogously.

The *m*_food_ and the *m*_sucrose_ are two free parameters. We fitted a single set of *m*_food_ and *m*_sucrose_ to all experimental values in Figure 5C and D, which is in total 8 points to fit (2 in panel C and 6 in panel D). The fitting was done by minimizing a least squares error between the model prediction and the corresponding experimental value. To identify the best fitting free parameters we re-parameterized them as *m*_food_ = *KR* and *m*_sucrose_ = *K*, where *R* is a the ratio of weights, and *K* is a scaling parameter. We sought *R* through grid searched between 1 and 10, with an interval of 0.1; then, a coefficient *K* was solved analytically for each *R* in the grid search to minimize a squared error between the model prediction and the corresponding experimental value. The above procedure of optimizing *R* and *K* is equivalent to optimizing *m*_food_ and *m*_sucrose_. The advantage is that if optimizing *m*_food_ and *m*_sucrose_, it has to be done via function evaluation (e.g., grid search on both of them); but while optimizing *R* and *K*, only *R* need to be optimized by function evaluation (in our case, grid search), but *K* can be solved analytically.

The experimental values were extracted from dopamine signals 2 seconds after the corresponding events as described in the caption of Figure 5. The extraction was done via a online digital plot extractor at https://apps.automeris.io/wpd/, our extraction can be loaded to this online two via “loading project”, and the project files is public here https://drive.google.com/drive/folders/1J4yQ3XjXebkNY2GNYrFttIfAT5ODuVJb?usp=sharing.

### Simulation of Dead Sea-salt experiment

In the simulated version of the experiments of Robinson and Berridge (2013) presented in Figure 6, the agents were exposed to the lever associated with the salt or the lever associated with the juice at random over 100 trials. If the salt lever was presented, the TD agent received a reward of −1 while if the juice lever was presented the agent received a value of +1. The TD agent learnt a value function with 2 states – juice and salt. The Reward Bases agent maintained separate salt and juice reward bases with the juice reward basis returning +1 for juice and 0 for salt and vice-versa for the salt basis. Reward Bases agent learned with state-modulated prediction errors. To match the rewards received, during training we set *m*_salt_ = −1 and *m*_juice_ = 1. The agents were trained with a learning rate of *α* = 0.1. Since there are no multi-step dependencies in this task, we trained with *γ* = 0.

To match the value functions of our agents to behavioural data, we assumed that the degree of appetitive approach and ‘liking’ behaviour plotted in the original experiment varied linearly to the value of the stimulus the animal had learned. That is, we simulated the simple equation,

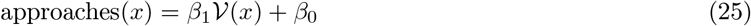

Regression parameters *β*_*j*_ were fitted for each model to data from the normal (non-depleted) condition. During testing in the deprivation conditions, the value functions remained the same for TD-learning, while they were recomputed for Reward Bases with *m*_salt_ = 0.5, as this value resulted in quantitative match with experimental data.

### Homeostatic Agent

Following Zhang et al. (2009), we implemented the ‘homeostatic agent’ which has also been proposed to explain instant generalization in reward revaluation experiments. The homeostatic agent uses the standard TD update rule, to learn a cached representation of the value function but also, at test time, can dynamically modulate its estimate of learnt values using a ‘homeostatic’ term *κ* which modulates the predicted reward (but not the final function). Specifically, at test time, the homeostatic agent estimates a saliency-modified value function 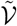 as,

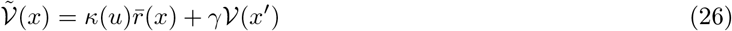

where *κ*(*u*) is a scalar parameter which is a function of the physiological state *u* and which scales the reward obtained in the value function update where 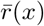 is the previously obtained reward in this contingency ^1^. The homeostatic agent can solve single-step tasks like the sea-salt experiment because it directly modulates the predicted reward from the outcome (salt or juice) according to the homeostatic state, however because it does not modulate the long-run value function, it cannot generalize to tasks with more than one step.

### Simulations of the Room Task

For all simulated agents, the value function was represented as a flattened vector of 6 × 6 = 36 states, which was initialized at 0. For the successor representation agent (see the next subsection for a detailed description of successor representations), the successor matrix was initialized as a 36 × 36 matrix of 0s and was updated on each timestep by the successor representation update rule (Equation 30). The successor agent computed its estimated value functions according to Equation 29.

For all agents a learning rate of *η* = 0.05, a discount factor of *γ* = 0.9 and a softmax temperature of 1 were used. Actions were selected by random sampling over the softmaxed distribution over actions. We used 500 steps between reversals. Means and standard deviations were obtained over 10 seeds for each agent in Figure 7.

### Successor Representation

The successor representation allows instant reward revaluation by computing and caching the “successor” matrix which is the matrix of expected state transition probabilities (Dayan, 1993). Given this matrix, it is possible to instantly recompute the value function if the reward function changes.

Let us first introduce notation required to describe the successor representation. Let **r** denotes a vector containing expected instantaneous rewards of all states in the environment, i.e. a vector of a length equal to the number of possible states, where each entry is equal to expected instantaneous reward for the corresponding state. Analogously, let **V** denote the vector of values of all states. Furthermore, let **𝒯** denote matrix with state transition probabilities, where entry **𝒯** _*y,x*_ denotes the probability of agent transitioning from state *x* to state *y* under the current policy. Using this notation, the definition of the value function from Equation 1 can be written as follows.

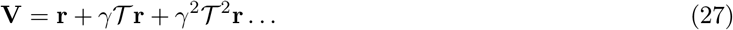

By a simple rebracketing, we can express this as,

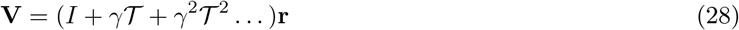

We can then separate out the matrix (*I* + *γ***𝒯** + *γ*^2^**𝒯** ^2^ …) and call it **ℳ**, the successor matrix giving us the following equation for the value function,

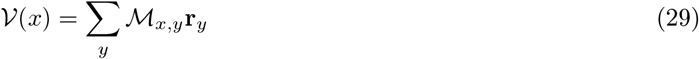

It is therefore clear that if we know **ℳ** and have it stored, then, given any change to the reward function **r**, it is trivial to recompute the correct value function. This allows for instantaneous reward revaluation since the value function can be recomputed so easily.

The successor matrix **ℳ** can be learned from agent’s experience. Each time an agent transitions from state *x* to *x′*, all entries in row *x* of matrix **ℳ** are updated,

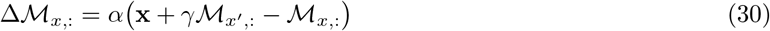

where **x** is one-hot state vector which signifies the state of the agent before the transition (i.e. its entry corresponding to *x* is equal to 1 while other entries to 0), and **ℳ**_*x*,:_ denotes the row *x* of matrix **ℳ**.

It is possible to show mathematically that the Reward Bases model is closely related to the successor representation and, indeed, can be intuitively thought of as a compressed successor representation with rewards tuned only to relevant dimensions. Recall that according to Equation 28, the value function can be expressed as **V** = **ℳr**, so analogously a value basis can be expressed as

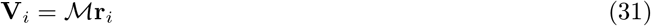

This can be demonstrated by substituting Equation 2 into Equation 28:

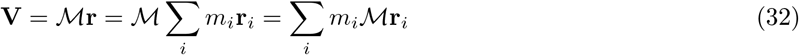

Since the value function also satisfies **V** = ∑_*i*_ *m*_*i*_**V**_*i*_, the condition of Equation 31 must hold.

The relationship of the Reward Bases model with the successor representation (Equation 31) helps explain why the Reward Bases model achieves equivalent performance to the successor representation at a significantly smaller computational cost.

## Acknowledgements

The authors would like to thank Nathaniel Daw and Scott Waddell for discussion and Mycah Banks for her aid in preparing the figures. This work has been supported by BBSRC grant BB/S006338/1 (B.M., Y.S., M.W. and R.B.), MRC grant MC_UU_00003/1 (R.B.), Wellcome Senior Research Fellowship 202831/Z/16/Z (M.E.W.), Henry Dale Fellowship from the Wellcome Trust (A.L.) and Royal Society 213465 (A.L.).

## Author contributions

Conceptualization, B.M. and R.B.; Software, B.M., Y.S. and R.B.; Formal Analysis, B.M. and A.L.; Data Curation, Y.S. and A.L.; Visualization, Y.S.; Writing - Original Draft, B.M. and R.B.; Writing - Review & Editing: Y.S., A.L. and M.E.W.

## Declaration of interests

The authors declare no competing interests.

## Data and Code Availability

At the time of publication, the data on dopaminergic responses of primate dopamine neurons analysed in this paper, and all codes will be made publicly available at Github.

## Footnotes

1 Another slightly different model also proposed by Zhang et al. (2009) is to define the reward term as log(*r*(*x*) + *κ*(*u*)). This approach only slightly changes the reward shaping and has no real impact on the fact that the homeostatic method, due to only modulating the reward, cannot perform zero-shot generalization across changing reward functions for multi-step tasks.

## Notes

### Competing Interest Statement

The authors have declared no competing interest.

